# KappaBle fluorescent reporter mice enable dynamic and low-background single-cell detection of NF-κB activity *in vivo*

**DOI:** 10.1101/2021.09.02.458704

**Authors:** Luigi Tortola, Franziska Ampenberger, Esther Rosenwald, Sebastian Heer, Thomas Rülicke, Jan Kisielow, Manfred Kopf

## Abstract

Nuclear factor-κB (NF-κB) is a transcription factor with a key role in a great variety of cellular processes from embryonic development to immunity, the outcome of which depends on the fine-tuning of NF-κB activity. The development of sensitive and faithful reporter systems to accurately monitor the activation status of this transcription factor is therefore desirable. To address this need, over the years a number of different approaches have been used to generate NF-κB reporter mice, which can be broadly subdivided into bioluminescence- and fluorescence-based systems. While the former enables whole-body visualization of the activation status of NF-κB, the latter have the potential to allow the analysis of NF-κB activity at single cell level. However, fluorescence-based reporters frequently show poor sensitivity and excessive background or are incompatible with high-throughput flow cytometric analysis. In this work we describe the generation and analysis of ROSA26 knockin NF-κB reporter (KappaBle) mice containing a destabilized EGFP, which showed sensitive, dynamic, and faithful monitoring of NF-κB activity at the single-cell level of various cell types during inflammatory and infectious diseases.

## Introduction

Nuclear factor-κB (NF-κB) encompasses a family of evolutionarily conserved transcription factors^1, 2^. Originally discovered as a transcriptional regulator for immunoglobulin κ-light chain ^3, 4^, NF-κB is now known to control a variety of cellular processes including proliferation, differentiation, apoptosis, and tissue homeostasis, in many cell types. In addition, NF-κB is of great importance in immunity and inflammation. It controls the transcriptional responses downstream of cytokine receptors and pattern recognition receptors as well as B- and T-cell receptors ^5^, and its deregulated activation is associated with the development of inflammatory pathologies.

The activity of NF-κB is regulated by members of a family of inhibitory subunits (IκB) that bind to it and prevent its nuclear translocation. Induction of NF-κB depends on the activation of upstream kinases of the IKK family that phosphorylate IκB, targeting it for ubiquitylation and proteasomal degradation. Upon release, NF-κB is free to migrate to the nucleus and act as a transcriptional activator. Two major branches of NF-κB signaling, namely the “canonical/classical” and “alternative” pathway can be distinguished. The canonical pathway relies on p65/p50 NF-κB heterodimers and is activated downstream of TLRs, as well as several cytokine receptors, including receptors of IL-1 family members and TNF. Whole body abrogation of the canonical pathway results in embryonic death caused by TNF-mediated liver wasting indicating its essential role in embryonic development ^6, 7^. While the canonical pathway is dispensable for lymphocyte development, it prevents TNF-driven apoptosis during inflammatory responses ^8–10^. Inversely, deregulated activation of the canonical pathway is associated with the development of immune pathologies ^11, 12^. The alternative pathway instead depends on RelB/p105 heterodimers and mediates signaling downstream of CD40 or LTβR, making it crucial for B cell responses and for the development of lymphoid structures ^13–17^.

Because of the prominent role of NF-κB in a broad variety of cellular processes, several approaches have been undertaken to monitor its activity by generation of reporter constructs. The majority of available reporters are based on three different approaches: 1) ectopic expression of a fluorescently-labelled NF-κB subunit and microscopy-based analysis of its cellular localization ^18^; 2) ectopic expression of a fluorescently-labelled IκB and detection of fluorescence loss during activation-driven IκB degradation^19^; 3) expression of a reporter protein (EGFP or luciferase) under the control of a NF-κB-dependent promoter ^20–23^. The first two systems have proven efficient in some settings, but their major drawback is the reliance on the (over)expression of fluorescently tagged components of NF-κB signaling, which may affect the functionality of the signaling protein and potentially alter cell physiology. These approaches also limit the analysis to a single member of the NF-κB (or IκB) family at a time, so that the dynamics of activation of other family members are inevitably overlooked. With respect to the first strategy, the necessity of determining the subcellular localization of fluorescent NF-kB subunits additionally precludes flow cytometric analysis. Systems relying on NF-κB-dependent expression of reporter genes such as fluorescent proteins or luciferase have the benefit of not affecting the upstream signaling pathway. The use of fluorescent reporters such as EGFP would furthermore enable unbiased, high-throughput, single cell analysis of heterogeneous cells mixtures by flow cytometry. However, the inherent stability and long half-life of these reporter proteins impedes a faithful analysis of the fine dynamics of NF-κB activation. Accordingly, existing EGFP expression-based reporter mice suffer from high fluorescence background and low sensitivity, which limits the usability of these systems ^22^. In this work, we tackle the inherent problems identified in previously generated reporter mice and describe the development of a low background, sensitive, fluorescence-based reporter system for the dynamic detection of NF-κB activity *in vivo* and the generation of ROSA26 knock-in “KappaBle” NF-κB reporter mice.

## Results

### Generation of a low-background fluorescent reporter for NF-κB using a self-inactivating retroviral vector

Aiming to generate a sensitive and low-background fluorescent reporter system for real-time analysis of NF-κB activity, we based our strategy on two main aspects (Fig. 1A): first, to increase the sensitivity of NF-κB reporter system ^24^, we used 8 NF-κB binding sites in the promoter region instead of 2-4 used in other reporter mice ^22, 23^; second, we used a destabilized version of EGFP containing a C-terminal PEST domain (destEGFP), which considerably reduces the half-life of EGFP and, hence, the background fluorescence in the absence of an activating stimulus. This should allow to better visualize the real-time dynamics of NF-κB activation ^25^.

**Figure 1.**
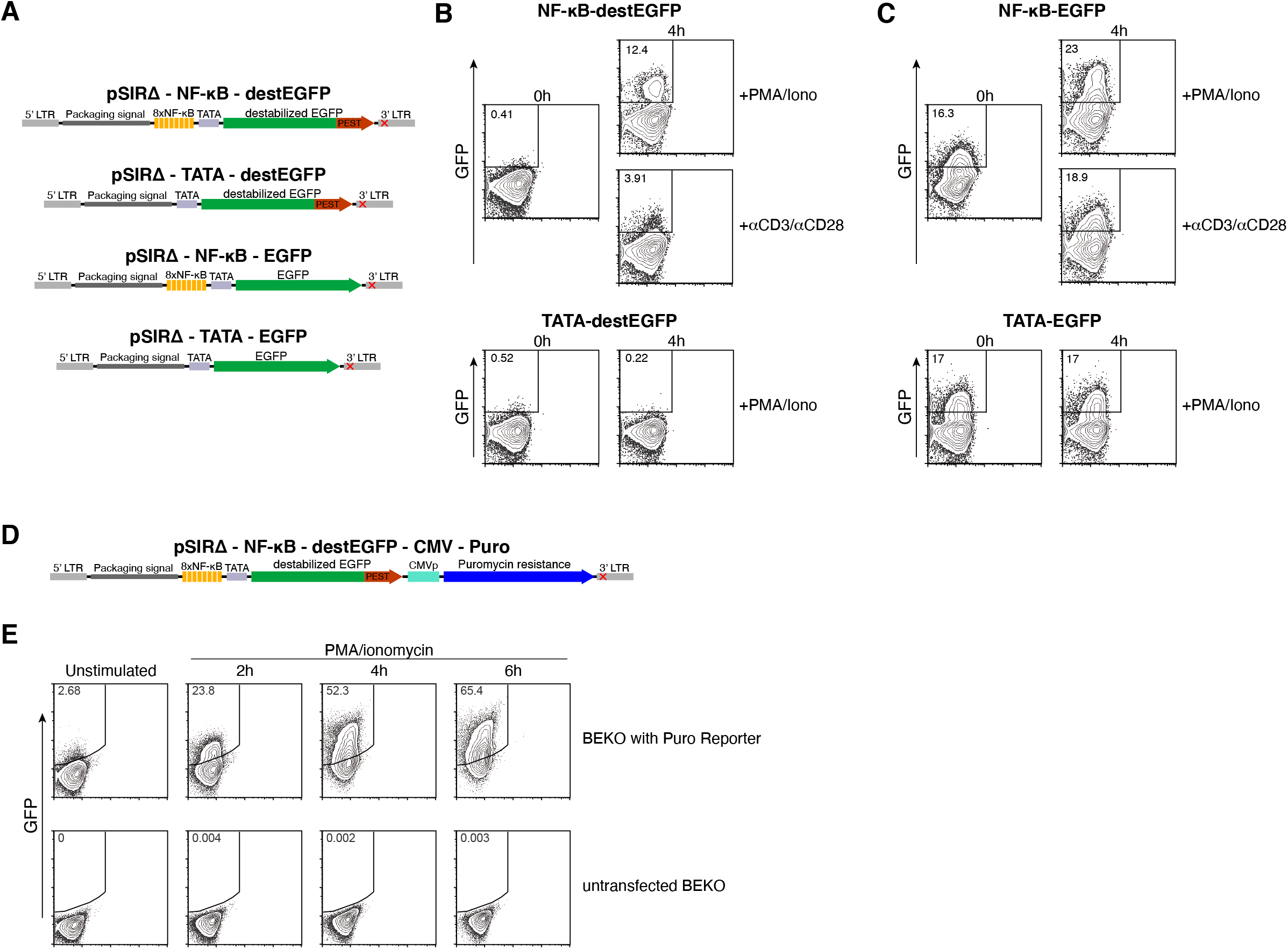
Sensitive, low-background detection of NF-κB activation with a retroviral reporter based on destabilized EGFP. (A) Schematic representation of pSIRΔ-NF-κB-destEGFP retroviral reporter vector and of the additional control constructs lacking NF-κB-binding sites and/or expressing conventional EGFP. LTR=long terminal repeat. The red cross in the 3’LTR indicates the mutation responsible for the inactivation of 5’LTR-dependent transcription upon integration into the host genome. (B) BEKO cells were infected with retroviral particles encoding NF-κB-destEGFP or TATA-destEGFP constructs. Cells were then stimulated with anti-CD3/anti-CD28 or with PMA and ionomycin and analyzed at the indicated time-points by flow cytometry to monitor GFP expression. (C) Flow cytometric analysis of BEKO cells carrying either the NF-κB-EGFP or the TATA-EGFP constructs after stimulation with PMA and ionomycin. (D) Schematic representation of pSIRΔ-NF-κB-destEGFP-CMV-Puro retroviral reporter vector. (E) BEKO cells were infected with retroviral particles encoding NF-κB-destEGFP-CMV-Puro. Following puromycin selection, cells were stimulated with PMA and ionomycin and analyzed at the indicated time-points by flow cytometry.

We incorporated the NF-κB-destEGFP construct into the backbone of a self-inactivating retrovirus (pSIRΔ). This vector enables delivery of the encoded reporter into dividing mouse cells and harbors a mutation that precludes 5’ long terminal repeat (LTR)-dependent expression of the reporter gene after integrating into the host genome ^26–28^. To highlight the advantage of destEGFP as a reporter protein, we additionally generated a retroviral NF-κB reporter using conventional EGFP (NF-κB-EGFP). Lastly, for both reporter versions, we generated additional control constructs lacking the NF-κB binding cassettes, while maintaining the TATA box. These controls (named “TATA-destEGFP” and “TATA-EGFP”) serve to distinguish the fluorescent background originating from steady-state or cumulative NF-κB activation over time from that dependent on TATA box-dependent basal transcription. All reporter constructs were then introduced and tested in the BEKO thymoma cell line ^29^. As shown in Fig. 1B, the NF-κB-destEGFP reporter led to virtually no fluorescence background in the absence of stimulation, while CD3/CD28 triggering or addition of PMA and ionomycin promptly induced EGFP expression in BEKO cells. Notably, a substantial and comparable proportion of EGFP^+^ cells was observed in BEKO cells containing the NF-κB-EGFP and TATA-EGFP in the absence of stimulation indicating a high fluorescence background driven by the TATA box, which could be increased by PMA/Ionomycin, but not CD3/CD28 stimulation, in an NF-κB-dependent manner.

This data indicates that usage of a destabilized EGFP is mandatory to faithfully monitor the dynamics of NF-κB activation, with a pronounced difference between stimulated and non-stimulated cells.

In order to allow the use of this reporter construct for applications that require the selection of transduced cells, we also generated an additional construct encoding puromycin-N-acetyl-transferase that confers resistance to puromycin (Fig. 1D). Stimulation of BEKO cells transduced with pSIRΔ-NFkB-destEGFP-CMV-Puro and subsequently selected with puromycin led to robust upregulation of GFP by the majority of cells (Fig. 1E). These data show that our retroviral destEGFP-based reporter shows minimal background fluorescence, allowing for a much more faithful detection of NF-κB activation compared to reporters based on conventional EGFP.

### Analysis of NF-κB activity in immune cells harboring the NF-κB-destEGFP reporter

Experiments performed in the mouse thymoma-derived BEKO cell line indicated that our destEGFP-based retroviral reporter could provide efficient and dynamic detection of NF-κB activation following stimuli such as CD3 crosslinking or mitogen-mediated activation. To determine the efficiency of this reporter in primary mouse cells, we infected mouse bone marrow cells with pSIRΔ-NF-κB-destEGFP. Interestingly, when culturing the bone marrow cells after retroviral infection, we observed that the addition of medium containing a fresh cytokine cocktail including IL-3, IL-6 and SCF led to an increase in the number of fluorescent cells and in the level of destEGFP expression as compared to cells cultured with cytokine-free medium (Fig. 2A,B). Importantly, once the cytokines were consumed by the BM cells, the residual fluorescence rapidly decreased, confirming the crucial advantage of using destabilized EGFP as a reporter over conventional EGFP for the observation of the dynamics of NF-κB activation.

**Figure 2.**
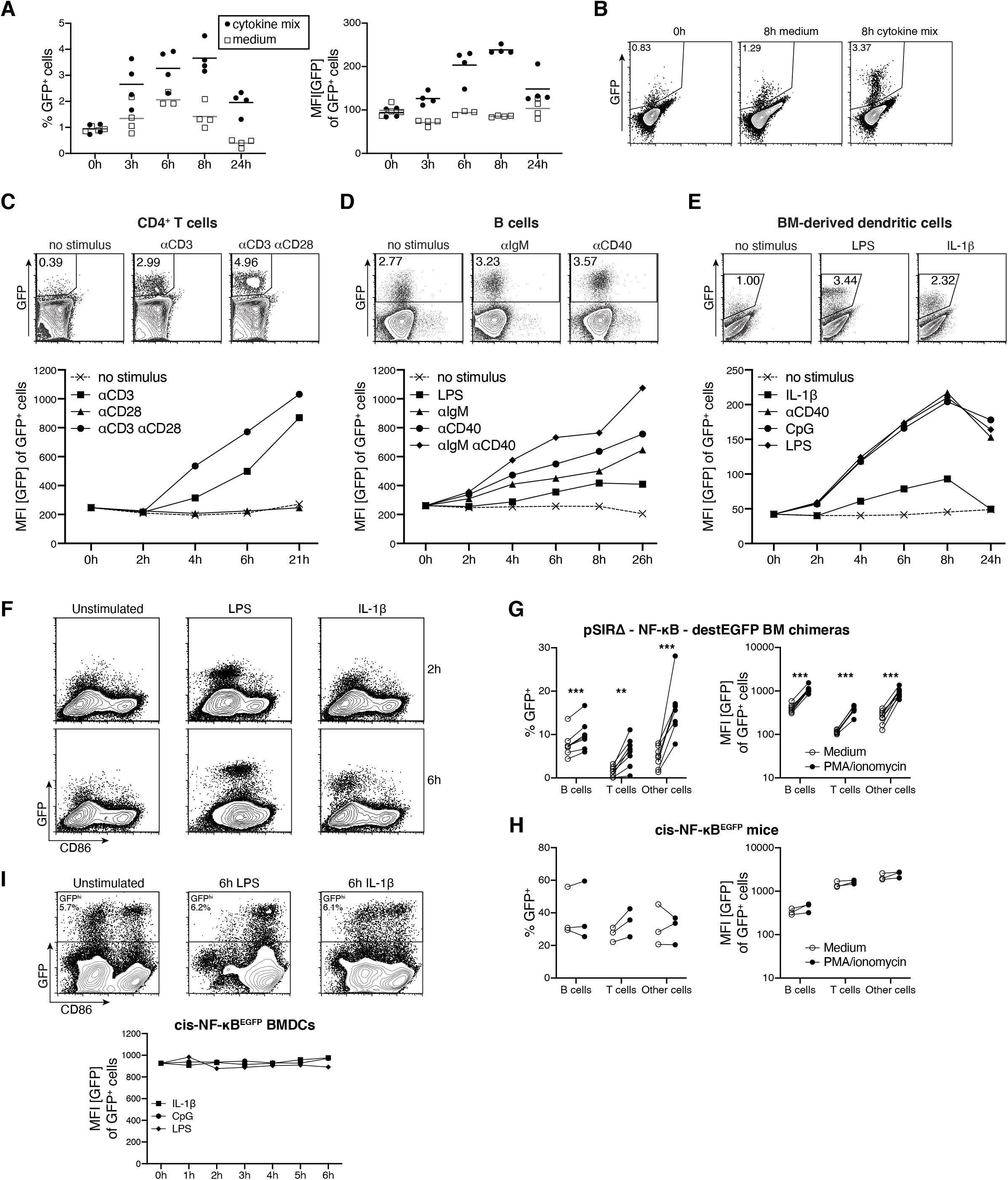
Faithful induction of destEGFP expression following activation of the canonical and alternative NF-κB pathways in primary immune cells. (A-B) BM cells infected with the retroviral NF-κB reporter were treated with a cytokine mix consisting of IL-3, IL-6 and SCF or with fresh medium without cytokines and analysed by flow cytometry at different time points. (A) Plots depict the percentage of GFP^+^ cells (left panel) and the MFI of GFP among GFP^+^ cells (right panel) at different timepoints after treatment. (B) Representative dot plots of the data shown in (A). (C-E) Lethally irradiated mice were reconstituted with BM cells infected with pSIRΔ-NF-κB-destEGFP retrovirus to generate NF-κB reporter chimeras. Following reconstitution, we isolated CD4+ T cells (C), B cells (D) and differentiated BMDCs (E), and treated them with a variety of stimuli to trigger NF-κB activation. Upper panels show representative contour plots from flow cytometric analysis and the lower panels show the mean fluorescent intensity (MFI) of GFP among GFP^+^ cells. (F) BMDCs derived from the BM of NF-κB reporter chimeras were left untreated or stimulated with LPS or IL-1β. Contour plots show the expression of destEGFP and CD86 at the indicated time points. (G-H) Blood cells isolated from retroviral NF-κB reporter chimeras (G) or from cis-NF-κB^EGFP^ mice (H) were left untreated or stimulated with PMA and ionomycin to assess the upregulation of GFP following NF-κB activation. Each circle indicates an individual mouse, paired observations from the same animal are connected with a line. Statistical analysis was performed using paired t tests with Holm-Sidak correction for multiple comparisons. (I) Bone marrow derived dendritic cells (BMDCs) from cis-NF-κB^EGFP^ mice untreated or treated with LPS or IL-1β. Contour plots on the top show the expression of destEGFP and CD86 at the indicated time points, whereas the plot at the bottom shows MFI of GFP among GFP^+^ cells.

Bone marrow cells transduced with pSIRΔ-NF-κB-destEGFP were then transplanted into lethally irradiated mice to generate NF-κB reporter chimeras. We then purified CD4^+^ T cells and B cells from the spleens of reporter chimeras by MACS and stimulated them *in vitro* to study the dynamics of destEGFP expression. Similar to BEKO cells, primary CD4^+^ T cells showed no expression of destEGFP (and hence no activation of NF-κB) in steady state. However, expression was rapidly induced upon CD3 cross-linking. Treatment with anti-CD28 antibodies alone did not lead to activation of NF-κB, but it enhanced destEGFP upregulation when combined with anti-CD3 antibody (Fig. 2C). B cells already expressed destEGFP before the addition of any exogenous stimulus, indicating that even at steady state B cells have a certain level of NF-κB activation, in agreement with previously published data ^4, 30, 31^. Importantly, stimulation of B cells with LPS, or by crosslinking IgM and/or CD40 on their surface, led to prompt upregulation of destEGFP expression as compared to non-stimulated cells (Fig. 2D).

To assess the functionality of the NF-κB reporter in myeloid cells, we differentiated BM-derived dendritic cells (BMDCs) from the bone marrow of reconstituted chimeric mice. We then stimulated BMDCs with LPS, CpG, anti-CD40 and IL-1β. DC activation led to destEGFP upregulation within 2 hours with TLR stimulation or CD40 crosslinking, or 4 hours when stimulating the cells with IL-1β(Fig. 2E). During the time-course, we also tracked the expression of the DC activation marker CD86 (Fig. 2F). Notably, destEGFP upregulation preceded the induction of CD86 following LPS treatment. As expected, in the case of IL-1β-driven NF-κB activation, destEGFP expression was induced in the absence of CD86 upregulation (Fig. 2F).

We next compared the sensitivity of our pSIRΔ-NF-κB-destEGFP reporter to a NF-κB fluorescent reporter mouse strain (i.e. cis-NF-κB^EGFP^) established previously ^22^. The reporter cassettes of the two systems differ in the number of NF-κB binding sites (8 vs 3) and in the reporter protein (destabilized vs conventional EGFP). Stimulation of blood cells isolated from pSIRΔ-NF-κB-destEGFP retro-transgenic mice with PMA and ionomycin led to a clear upregulation of GFP (Fig. 2G) on B cells, T cells, and myeloid cells. Blood cells from cis-NF-κB^EGFP^ animals showed high background fluorescence in the absence of any stimulus, and *in vitro* stimulation with PMA and ionomycin did not result in further upregulation of GFP (Fig. 2H). Similarly, stimulation of cis-NF-κB^EGFP^ BMDCs with LPS or IL-1β did not upregulate GFP above levels of unstimulated cells (Fig 2I).

Collectively, these data demonstrate that the retroviral destEGFP-based reporter can be employed to visualize the activity of NF-κB in primary murine immune cells *ex vivo* by flow cytometry. The elevated number of NF-κB response elements in the promoter region and the use of destabilized EGFP as a reporter protein leads to a significant increase in sensitivity compared to previously generated fluorescent reporter animals. Furthermore, we also show that our reporter can successfully portray the activation of both major pathways of NF-κB signaling, i.e. the classical (for TLRs, IL-1R, BCR and TCR) and the alternative pathway (CD40).

### Generation of KappaBle NF-κB reporter mice and monitoring of NF-κB activity in a variety of cells in the lung during inflammation

The results presented in the previous sections show that our construct allows sensitive, quantitative and rapid detection of NF-κB activation in hematopoietic cells. Retro-transgenesis is a suitable solution to incorporate our reporter in immune cells, but in order to be able to track NF-κB activity across all cell lineages, we decided to generate knock-in mice encoding the reporter construct in the ROSA26 locus ^32^. After transfection of C57BL/6-derived ES cells and geneticin selection, integration of our targeting construct by homologous recombination was assessed by PCR on the short arm (Fig. 3A). Successfully targeted ES cells were used to generate “KappaBle” NF-κB reporter mice. Lymphocytes isolated from the blood of KappaBle mice (Fig. 3B) lacked background fluorescence, and stimulation with PMA and ionomycin lead to a significant and faithful upregulation of GFP, although only in a relatively small fraction of T cells (~3%) and B cells (~0.4%). Hetero- or homozygous expression of the KappaBle cassette yielded comparable results (data not shown). On the other hand, intratracheal administration of LPS led to striking upregulation of GFP in alveolar macrophages (AMs, >70%), DCs (~15%) and CD31^+^ endothelial cells (~40%) in the lungs 5 hours post injection (Fig. 3C). Importantly, this GFP signal significantly faded 18 hours after the administration, highlighting again the advantage of destEGFP as a reporter protein to enable dynamic measurement of NF-κB activity. Considering that a large fraction (>65%) of BEKO thymoma cells containing pSIRΔ-NFkB-destEGFP potently upregulate GFP expression upon stimulation, we were wondering why only a small fraction of T cells (and B cells) from KappaBle mice upregulate GFP upon PMA/Ionomycin stimulation. To address this question, we generated KappaBle T cell hybridomas by fusion of T cells from the spleen of KappaBle mice with BW5147 cells. Interestingly, the resulting clones showed enormous range of maximal GFP responses following PMA/ionomycin stimulation, ranging from ~0.5 to >80% (Fig. 3D), indicating that random epigenetic changes associated with the fusion determine the accessibility of the reporter cassette to the transcriptional machinery. Nonetheless, background fluorescent in the absence of a stimulus was consistently extremely low across all T cell hybridoma clones, further confirming the specificity and low background of our reporter system.

**Figure 3.**
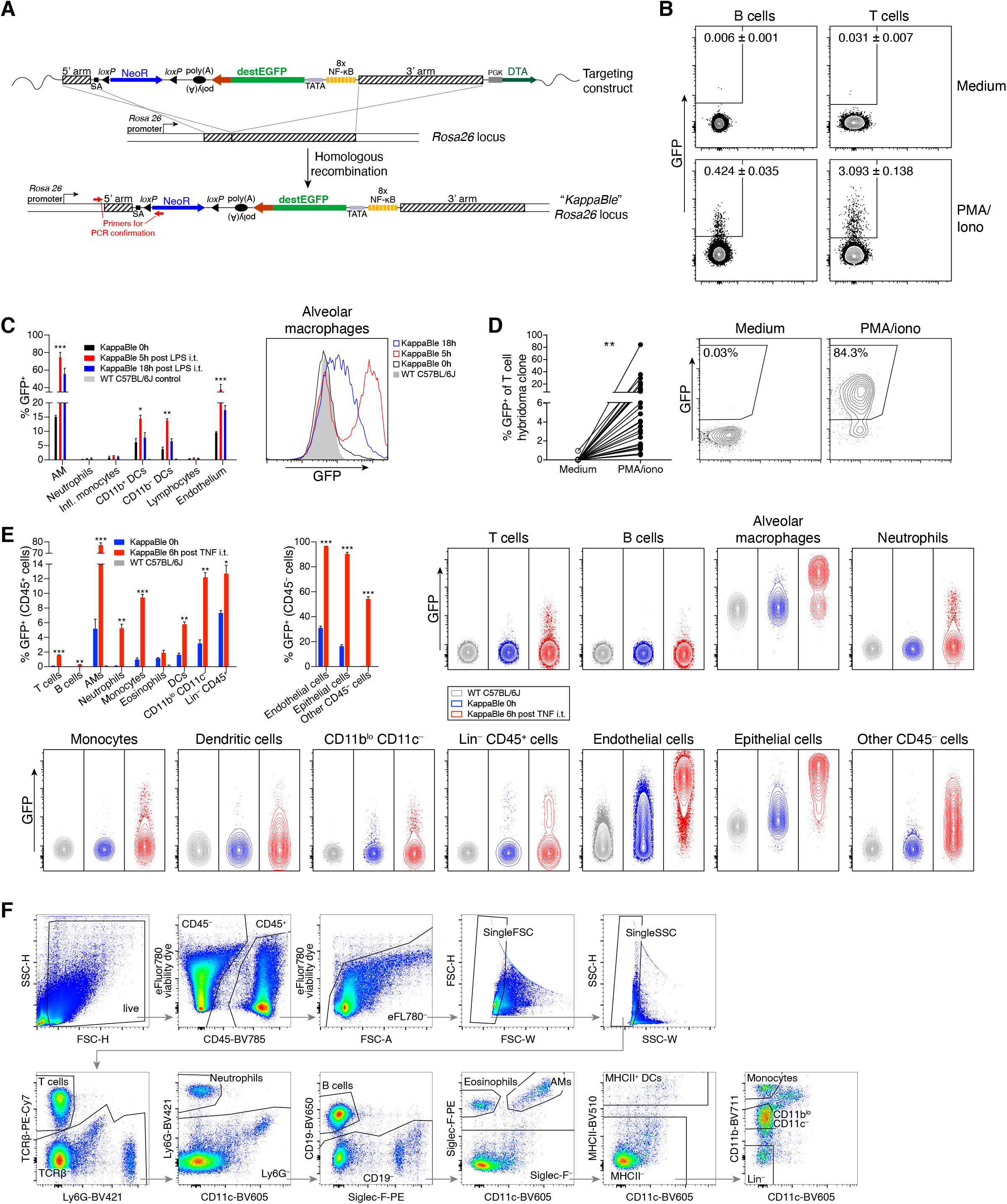
Generation and characterization of KappaBle NF-κB reporter mice. (A) Schematic representation of the vector generated to deliver the NF-κB reporter construct to the ROSA26 locus in ES cells and of the targeted “KappaBle” locus after homologous recombination. (B) Cytometry plots show destEGFP expression in B and T cells from the blood of KappaBle mice (n=9) before and after *in vitro* stimulation with PMA and ionomycin. (C) KappaBle (C57BL/6J) and C57BL/6J control mice were treated with LPS intratracheally and analyzed by flow cytometry at the indicated time points after injection. Bar graph on the left shows % of GFP^+^ cells in the indicated populations and timepoints and histogram overlay on the right shows GFP expression in AMs at different timepoints. Statistical analysis was performed using 2-way ANOVA with Dunnett correction for multiple comparisons (3 mice per group/timepoint). (D) Hybridomas were derived from KappaBle T cells. Individual hybridoma clones (n=26) were left untreated or stimulated with PMA and ionomycin. Cells were then analyzed by flow cytometry. GFP expression is plotted on the left panel, paired observations from each individual clone are connected by a line (statistical analysis was performed with paired t test). The contour plot on the right shows expression of GFP^+^ by the most responsive KappaBle T cell hybridoma clone. (E) KappaBle and WT C57BL/6J mice were treated with TNF intratracheally and analyzed by flow cytometry 6 hours after injection. Bar graphs on the top left show the percentages of GFP^+^ cells among indicated populations of CD45^+^ and CD45^−^ cells, while contour plots on the right depict GFP expression in the indicated populations and groups. Statistical analysis was performed using paired t tests with Holm-Sidak correction for multiple comparisons (3 mice per group/timepoint). (F) Gating strategy for the flow cytometry data depicted in (E).

Receptors for TNF (TNFR1 and/or TNFR2) are broadly expressed by many cell types and are known to trigger NF-κB activation after engagement with their ligand. To visualize NF-κB activity across different cell types in an inflammatory setting, we administered TNF by intratracheal instillation to the lungs of KappaBle mice and analyzed a variety of hematopoietic and non-hematopoietic cells isolated from the lungs by flow cytometry 6 hours later (Fig. 3E,F). Similar to endotoxin exposure, we found a strong responsiveness of AM to TNF (>70% GFP^+^). Other immune cell types, including neutrophils, monocytes, DCs, CD11b^lo^ CD11c^−^ cells (predominantly NK cells) and Lin^−^ cells (including γδ T cells and ILCs) showed a sizable upregulation of GFP (5-15%) following exposure to TNF *in vivo*. B and T cells isolated from the lungs of KappaBle mice showed measurable and consistent upregulation of GFP after NF-kB activation, but only in a minority of cells, consistent with the observations in the blood. Eosinophils were ~1% GFP^+^ in untreated mice, and this percentage slightly increased following TNF administration.

Importantly, analysis of the non-immune CD45^−^ compartment in the lungs revealed that TNF triggered the upregulation of GFP in almost all EpCAM^+^ epithelial and CD31^+^ endothelial cells, as well as in ~50% of other CD45^−^ stromal cells.

These data show that the NF-κB reporter in KappaBle animals displays minimal background and is functional across all immune cell types we analyzed, although the percentage of responding cells varies. Among CD45^−^ non-immune cells this percentage greatly increased and almost all epithelial and endothelial cells responded to NF-κB activation by upregulating GFP.

### Influenza-infected alveolar macrophages and lung epithelial cells show increased NF-κB activation

We next studied the dynamics of NF-κB activation in the lungs of influenza-infected KappaBle mice. Analysis of immune cells 5 days post-infection showed a clear upregulation of GFP in AMs and γδ T cells compared to naïve mice (Fig. 4A,B). Among non-immune cells, both epithelial and endothelial cells showed significantly increased GFP expression following infection (Fig. 4A,B). We then wondered whether NF-κB activation correlated with whether a cell had been infected or not. Intracellular staining of influenza nucleoprotein (NP) showed that the virus predominantly infected (or was taken up by) AMs and neutrophils and a small proportion of epithelial cells (Fig. 4C). Interestingly, NP staining in sorted GFP^+^ and GFP^−^ cells showed that infected cells were significantly enriched among GFP^+^ AMs and GFP^+^ epithelial cells (Fig. 4D), although not every infected AM and epithelial cell was also GFP^+^, indicating that infection did not necessarily activate NF-κB. Neutrophils showed a similar, albeit statistically non-significant, trend. Moreover, although the frequency of GFP^+^ endothelial cells and γδ T cells increased during infection, NF-κB activation in these populations seems to be driven by inflammation rather than direct viral infection since they were negative for NP. These data highlight the potential of KappaBle mice for the unbiased analysis of NF-κB activity during complex cellular responses *in vivo*.

**Figure 4.**
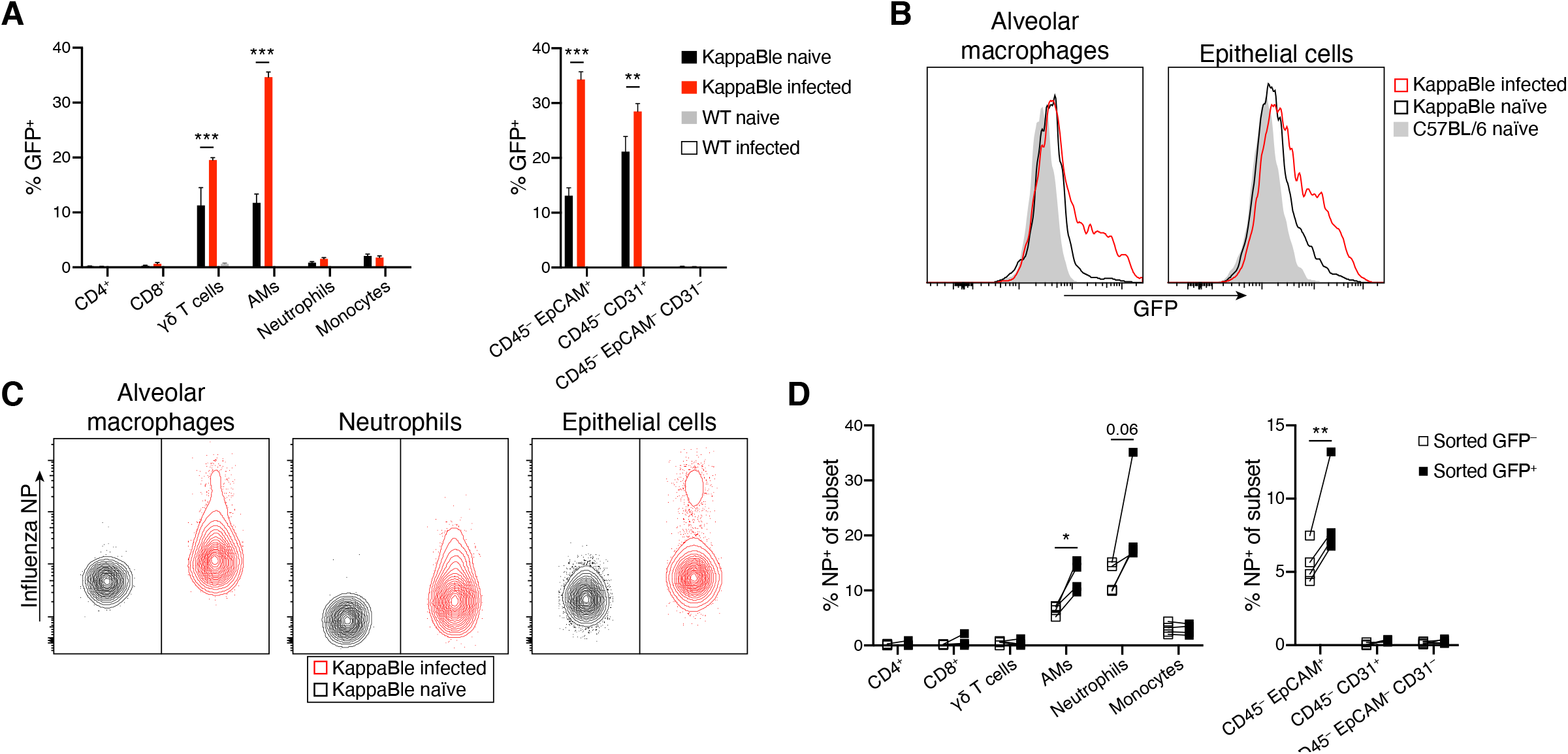
Analysis of NF-κB activity in the lungs of influenza-infected KappaBle mice. (A-D) KappaBle mice (n=4) were infected with influenza A virus. On day 5 post infection lung cells were isolated and analysed by flow cytometry. (A) Bar graphs show GFP expression in the indicated populations of CD45^+^ and CD45^−^ cells. Statistical analysis was performed using 2-way ANOVA with Sidak correction for multiple comparisons. (B) Representative histogram overlays depict GFP expression in AMs and lung epithelial cells. (C) Lung cells were stained intracellularly for influenza NP to detect infected cells. Contour plots depict representative NP staining in AMs, neutrophils and lung epithelial cells. (D) NP staining was performed in sorted GFP^+^ and GFP^−^ cells from infected mice. Plots depict % NP^+^ among CD45^+^ (left panel) and CD45^−^ cell populations (right panel). Each symbol represents an individual mouse, paired observations from the same mouse are connected with a line. Statistical analysis was performed using paired t tests with Holm-Sidak correction for multiple comparisons.

## Discussion

NF-κB plays an essential role in a plethora of cellular processes. In order to efficiently track the activation of this transcription factor, we set out to generate a reporter construct based on GFP expression, to allow for rapid, high-throughput analysis by flow cytometry. Common problems encountered when using GFP as a reporter gene are the low sensitivity and the high levels of background fluorescence ^33^. In order to overcome such obstacles, we generated a retroviral vector encoding a destabilized EGFP version with a short half-life under the control of multiple NF-κB binding sites. Expression of this vector in a thymoma cell line and primary mouse immune cells showed inducible, faithful and dynamic expression of GFP when activating NF-κB with various stimuli, such as TLR ligands, CD3-crosslinking, or cytokines, and a low fluorescent background in the absence of a stimulus. Importantly, due to the overlapping reactivity of the response element with the various isoforms of NF-κB, we could efficiently detect activation of both the canonical pathway, induced for instance by TLR ligands or IL-1β, and of the alternative pathway, triggered by CD40 crosslinking, making our retroviral NF-κB reporter a powerful tool for the detection of NF-κB activation.

In order to allow the measurement of the dynamics of NF-κB activation also in non-hematopoietic cell lineages *in vivo*, we generated KappaBle NF-κB reporter mice. We decided against random transgenesis to avoid adverse integration effects and chose to introduce the reporter into the ROSA26 locus, which has been previously described to be accessible in the vast majority of cell types ^34^. Analysis of hematopoietic cells from KappaBle mice showed that the reporter cassette was functional across all analyzed immune cell types and *in vivo* or *in vitro* stimulation with NF-κB-activating stimuli led to consistent upregulation of GFP. However, the fraction of cells responding to NF-κB activation with the expression of GFP varied in the different cell types: after *in vivo* stimulation, lung resident AMs showed an almost universal response, while other myeloid cells (including DCs, neutrophils and monocytes) showed upregulation of GFP in a subset of cells (5-15%). Among lymphocytes, NK cells and γδ T cells/ILCs showed the most pronounced upregulation of GFP upon activation (>10%), while the fraction of GFP-expressing cells was relatively low among αβ T cells (~3%) and, in particular, B cells (<1%). This could be explained by epigenetic silencing of the reporter construct or expression of a repressor that binds to the NF-κB motif and prevents transcription in certain cell types. This hypothesis is supported by the analysis of hybridomas derived from KappaBle T cells, in which the maximal upregulation of GFP greatly varied among different clones.

Importantly, analysis of the non-immune compartment of KappaBle mice following *in vivo* NF-κB activation showed GFP upregulation by all lung epithelial and endothelial cells and by ~50% of remaining stromal cells, demonstrating the great potential of KappaBle mice for the study of NF-κB activity in non-immune cells *in vivo*. To highlight the potential application of our reporter animals for the analysis of NF-κB in complex *in vivo* cellular processes, we investigated the activity of NF-κB in the lungs of influenza-infected KappaBle mice and found that it was primarily activated in virus-infected AMs and lung epithelial cells.

In summary, we describe here the generation of a fluorescence-based, low-background reporter system for the study of NF-κB. Both the retroviral reporter and knock-in KappaBle reporter mice offer the possibility to track the dynamics of NF-κB activity during complex processes *in vivo* or *in vitro* at the single cell level, making them useful tools for researchers studying the role of NF-κB in health and disease.

## Materials and methods

### Cloning of retroviral NF-κB reporter vector and of KappaBle targeting construct

The octameric NF-κB response element was generated by multimerizing a sequence encoding two distinct NF-κB binding sequences ^24^, which was obtained by annealing of two synthetic oligonucleotides with complementary 5’ overhangs (sequences: 5’-agc tTG GGG ACT TTC CGC GAG GGA TTT CAC CTA-3’ and 5’-agc tTA GGT GAA ATC CCT CGC GGA AAG TCC CCA-3). The octamer was gel-purified, amplified by PCR with Platinum Taq polymerase (Invitrogen) using single stranded oligonucleotides as primers and finally re-purified from the gel. The product was cloned into p123TA vector and sequenced. The octamer was then extracted from the vector by BamHI (made blunt) and EcoRI digestion and cloned into pBluescript II-KS(+) digested with EcoRV and EcoRI to generate pBS NFκB. The IL-2 minimal promoter sequence containing a TATA-box was amplified from mouse genomic DNA with Pwo polymerase (synthesized and purified in-house as previously described ^35^) using the primers 5’-agt att ggt ctc gaa tt g CAT CGT GAC ACC CCC ATA TTA T-3’ and 5’-agg gat ccA GGC AGC TCT TCA GCA TGG GAG-3’ ^24^. The PCR product was digested with BsaI and EcoRI and cloned into pBS-NFκB cut with BamHI and EcoRI to yield pBS-NFκB-IL2mp. Destabilized EGFP (destEGFP) was extracted from the pd2EGFP vector and cloned into pBS-NFkB-IL2mp by using NotI and BamHI, generating pBS-NF-κB-destEGFP. The construct was sequenced and the NF-κB reporter cassette was transferred into pSIRΔ vector (Clontech) using XhoI and NotI. To generate the reporter construct carrying puromycin resistance (Fig. S1A,B), the cassette from pSIRΔ-NFkB-destEGFP was cloned into pSIRΔ-U6-CMV-Puro using XhoI and BglII.

To generate the KappaBle embryonic stem (ES) cell targeting construct, a synthetic double-stranded oligonucleotide containing a loxP site was first cloned into the pBS NF-κB reporter vector using SacII and NotI. The sequence of the oligonucleotide was 5’-TAC CGC GGT CTC GAG GTC ATG CAT CAT AAC TTC GTA TAA TGT ATG CTA TAC GAA GTT ATT AAT AAA GAC GCG TCT CTA GAG CGG CCG CTA-3’. Following this step, a 240-bp fragment from pHL-HH vector (courtesy of H. Fehling ^36^) encoding additional polyadenylation signals was cloned using MluI and XbaI, yielding pBS NF-κB loxP polyA. The long arm of homology from pHL-HH was cut with Asp718 and AscI (blunt) and cloned into pBS NF-κB loxP polyA opened with Asp718 and SalI (blunt), generating pBS NF-κB loxP polyA LA. The reporter-LA cassette was then excised and cloned into pHL-HH by using NsiI and Asp718, to yield the final KappaBle targeting vector. All the enzymes used for restriction digestion were purchased from New England Biolabs or from Roche Applied Science.

### Mice

Mice with specific pathogen-free status according to FELASA were bred and maintained in individually ventilated cages at the ETH Phenomics Facility (Zurich, Switzerland). For experiments, age-matched mice in the age of 6-10 weeks were used. Swiss federal and local animal ethics committees approved the described animal experiments.

### Cell preparation

Single cell suspensions from the spleen were prepared by pressing the organ through 70μm pore size strainers (BD Biosciences) in PBS 2% FCS. To obtain splenic dendritic cells, spleens were minced and digested 45 minutes with collagenase IV (300units/mL) and DNaseI (200units/mL, both enzymes from Worthington) before filtration through cell strainers. To isolate lung cells, IMDM medium supplemented with collagenase IV (600units/mL) and DNaseI (200units/mL) was introduced into the lungs of euthanized mice by cannulating the trachea. Perfused lungs were then digested for 45 minutes under agitation at 37°C, followed by fine mincing and filtration through 70μm cell strainers. For the analysis of intratracheally-treated TNF-treated mice, lungs were digested using Liberase TM (Sigma, 50 μg/mL) and DNaseI (200units/mL) and processed using the gentleMACS dissociator (Miltenyi). For some experiments, CD4^+^ T cells and B cells were MACS-sorted with anti-CD4 and anti-CD19 microbeads (Miltenyi), respectively. For the generation of bone marrow-derived dendritic cells (BMDCs), bone marrow (BM) cells were obtained from donor mice by flushing the femurs and the tibiae with PBS 2% FCS and cultured on bacterial, non-treated Petri dishes in RPMI medium supplemented with 10% FCS, HEPES, glutamine, penicillin/streptomycin and granulocyte-monocyte colony stimulating factor (GM-CSF, supernatant from X63-GMCSF cell line). Fresh medium was added on day 3, 6 and 8. For experiments, BMDCs were collected on day 9 by gentle pipetting from the culture plate.

### Retroviral transfection

Retroviral particles were assembled using the Phoenix packaging cell line ^37^. Cells were seeded in complete IMDM (10% FCS, P/S, β-ME). About 5 min prior transfection, fresh medium containing 25 μM chloroquine was given to the cells. Separately, 50 μg of the retroviral construct were resuspended in 1752 μl of ddH_2_O, and 248 μl of a 2M CaCl_2_ solution were added. After that, 2 ml of 2x HBS solution (50mM HEPES, 10mM KCl, 12mM dextrose, 280 mM NaCl, 1.5mM Na_2_HPO_4_, pH 7 ± 0.1) were added with a 5 ml pipette while vigorously aerating. This mix was then immediately added to the cells, which were then incubated for 8h. After this time, fresh medium was added to the cells and the retroviral supernatant was collected 24h later. This procedure was repeated once more before discarding the cells. Supernatant was stored at −80°C till use. For retroviral transfection of bone marrow cells, donor mice were injected with 5-fluorouracil (150 μg/g bodyweight) intraperitoneally. Six days later, donor mice were sacrificed, the hind limbs removed and the bone marrow collected by flushing the femurs and the tibiae. Cells were cultured in a 24-well plate (500’000 cells/well) in complete IMDM supplemented with 6 ng/ml IL-6 (Sigma), 6 ng/ml IL-3 (eBioscience) and 10ng/ml SCF (produced in-house). On the following day, retroviral supernatants were supplemented with 4μg/ml polybrene and the cytokines from above, and distributed on the cultured BM cells. Spin infection was performed by centrifuging cells at 1800 rpm for 45 min, incubation for 1h at 37°C and repetition of the centrifugation after providing fresh retroviral soup. The whole procedure was then repeated the following day, before reconstitution of lethally irradiated recipient mice with at least 5×10^5^ transfected BM cells. For transfection of BEKO cells ^29^, minor modifications of the protocol from above were used. No cytokines were supplemented in the medium and only one spin infection round (1800 rpm for 30 min) was performed.

### In vitro stimulations

For the experiments described in the manuscript, soluble stimuli were used at the following concentrations: lipopolysaccharide (LPS, InvivoGen) 500ng/ml, CpG (InvivoGen) 100nM, IL-1ß (ThermoFisher) 200ng/ml, anti-CD40 (clone 1C10, Biolegend) 1μg/ml, PMA (Sigma) 10^−7^M, ionomycin (Sigma) 1μg/ml. For plate coating, anti-CD3ε (clone 145-2C11, Biolegend) and anti-CD28 (clone 37.51, Biolegend) were used at a concentration of 2μg/ml, at and anti-IgM F(ab’)_2_ (Jackson Immunoresearch Laboratories) at 10μg/ml.

### In vivo treatments and infections

For LPS treatment, 100ng LPS suspended in 50μL PBS were injected intratracheally and animals were analysed 5 and 18 hours post injection. TNF (Peprotech) was also administered intratracheally (1μg in 50μL PBS) and animals were analysed 6 hours later. Influenza infection was performed by intratracheal administration of 500pfu Influenza virus strain PR8 (A/Puerto Rico/34, H1N1). On day 5 post-infection, animals were euthanized and lung cells were isolated for analysis.

### ES cell culture and selection and blastocyst injection

LK1 C57BL/6J mouse ES cells ^38^ were grown on a layer of irradiated neomycin-resistant EFs in ES medium (DMEM Glutamax supplemented with 1’000 units/ml LIF, 15% ES grade FCS, 1mM sodium pyruvate, MEM non-essential amino acids, penicillin/streptomycin, β-mercaptoethanol). On the day of transfection, ES cells were trypsinized and washed in PBS. 10^7^ cells ES cells were suspended in 800μl transfection mix (HEPES 20 mM, pH7.0, NaCl 137 mM, KCl 5 mM, Na_2_HPO_4_ 0.7 mM, Glucose 6 mM, β-Mercaptoethanol 0.1 mM) containing 30 μg KappaBle targeting construct which had been linearized overnight with PvuI and purified by ethanol precipitation. This mix was then transferred to an electroporation cuvette (0.4cm) and electroporated with 0.240 kV, 475 μF. Cells were incubated 10 minutes at room temperature, then transferred in ES medium and distributed on 3 culture dishes. Geneticin (175 μg/ml) was added for selection starting 1 day after transfection. Medium was changed daily until day 7, when colonies were picked and cultured in 96-well plates. Screening for homologous integration of the short arm was performed by a 2-round nested PCR on the ES DNA. The primers used for the first round were 5’-CCT AAA GAA GAG GCT GTG CT-3’ (ExtRnd1), 5’-CAT CAC AGC AGC CGA TTG TC-3’ (NeoRnd1), whereas those for the second round were 5’-AGA GAG CCT CGG CTA GGT AG-3’ (ExtRnd2), 5’-CAT AGC CGA ATA GCC TCT CC-3’ (NeoRnd2). The reaction was carried out using Platinum Taq polymerase, in the presence of 5% DMSO. Each round of PCR consisted of 30 cycles of 95°C for 30 sec, 56.5 for 30 sec, 68°C for 3 min. Of 36 picked clones, 6 were found to be positive and were further expanded and subsequently frozen in ES freezing medium (90% ES medium, 10% DMSO). 3 positive clones were karyotyped, microinjected into BALB/c blastocysts and transferred into the uterus horns of pseudopregnant surrogate mice. Chimeras were obtained and crossed to C57BL/6J mice. Twenty-one black offspring were screened for the presence of the neomycin resistance cassette, 9 were found to be positive and named “KappaBle” mice.

### T cell hybridoma generation

Splenic T cells isolated from KappaBle mice were activated with plastic-bound anti-CD3ε and anti-CD28 Abs in the presence of mouse IL-2 for 2-3 days. Equal numbers of T cells and BW5147 cells were then fused using PEG-1500 and plated at limiting dilution in the presence of 100mM hypoxanthine, 400nM aminopterin, and 16mM thymidine (HAT) and after selection single clones were isolated.

### Flow cytometry and data analysis

For surface staining, cells were resuspended in PBS 2% FCS, briefly incubated with Fc receptor blocking mAb (clone 2.4G2) and then with fluorescent labelled surface antibodies at 4°C for 15 minutes. For intracellular staining, surface-stained cells were fixed with 4% formalin (10 minutes) and permeabilized with PBS 2% FCS 0.5% saponin. Cells were then incubated with fluorescently labelled antibodies for 30 minutes (room temperature). Influenza nucleoprotein (NP) was stained using clone HB65 (homemade), followed by secondary staining with anti-mouse IgG-PE. Fluorescent labelled antibodies were purchased from Biolegend, ThermoFisher and BD Biosciences. After staining, cells were washed and resuspended in PBS 2% FCS for flow cytometric analysis and sorting using a FACSCalibur, a FACSCanto, a LSRFortessa and a FACSAria Fusion (all instruments from BD Biosciences). FACS data was then analyzed using FlowJo and statistical analysis was performed using Prism (Graphpad software). Throughout the article, (*) indicates a P-value <.05, (**) a P-value of <.01 and (***) a P-value of <.001.

## Acknowledgements

We thank the teams of the ETH Flow Cytometry Core Facility for cell sorting and the ETH Phenomics Center (EPIC) mouse facility for animal husbandry. Peter Nielsen for discussions and reading of the manuscript. We are grateful to the Swiss National Science Foundation (SNF) for funding (grants 310030_163443 and 310030B_182829).

## Author contributions

LT, JK, and MK conceived the study; LT and JK initiated study design; LT performed majority of experiments; FA, ER, and SH contributed to data acquisition; TR secured blastocyst injection of ES cells and generation of the mice; LT drafted manuscript; MK critically reviewed and revised manuscript and secured funding of the study.

